# Microbes under climatic refugia: Self-stabilizing subcommunity rank dynamics in large-river deltaic estuaries

**DOI:** 10.1101/2025.01.25.630657

**Authors:** Huaiqiang Liu, Zhenghua Zhong, Juan Xu, Liping Qu, Haijun Liu, Fengfeng Zheng, Chuanlun Zhang, Wei Xie

## Abstract

Microorganisms, akin to macroorganisms, can be sheltered by climatic refugia, overlooked for optimism about microbial extinction. By categorizing microbial communities into ecologically distinct subcommunities, we conducted a two-year survey of large-river deltaic estuaries (LDEs) which are regarded as refugial hotspots, and established rank hierarchies to monitor temporal changes. We found that, consistent with macroecology, LDEs harbor distinct microbial community structures, with the abundance distribution of subcommunities exhibiting resilience against environmental perturbations, persisting on an annual cycle. In particular, while the evenness of subcommunity rank abundance curves fluctuates frequently, shifts in rank order are less dynamic but, once initiated, have a more pronounced impact on subsequent rank dynamics. Our results suggest that the microbial subcommunity rank structure in LDEs is both persistent and robust, aligning with classic patterns of species abundance distributions. This study emphasizes that, despite differences in their roles, the stable functioning of microbial rank systems is also fundamental to the refugial capacity of ecosystems and should be given equal importance.

## Introduction

The impact of climate fluctuations on populations may not always be adverse, particularly for those that are ephemeral but not at risk of extinction (MacDonald et al., 2024). A more common option, however, is to move to refugia which can buffer against climate change (Morelli et al., 2016). Unlike glacial refugia during the period of glaciation, which are caused by the glacial and interglacial expansion–contraction cycle (Stewart et al., 2010), climatic refugia are more generalized and can be biogenic (Jurgens & Gaylord, 2018), topoclimate-shaped (Ackerly et al., 2020) or even ecosystem-protected (Morelli et al., 2020). Climatic refugia can also be created by food supplies. For instance, the presence of glacial plumes in front of tidewater glaciers generates a climatic refugium that increases prey availability (Hop et al., 2023). Furthermore, the protection of central, palatable seedlings from herbivores by the presence of surrounding, undesirable neighbors can also generate an associational refugium (Finnerty et al., 2024). Refugia are occasionally triggered only during particular episodes, such as thermal refugia for hot weather (Sullivan et al., 2021; Valentine et al., 2024) and flow refugia for flood events (Fuller et al., 2010; Mathers et al., 2022).

Amidst the variety of prefixes, geographical descriptors consistently emerge as the most prevalent. Montane environments, rocky environments, and riparian/terrestrial wet spots are three recognized types of terrestrial refugia, with the latter least well-documented (Selwood & Zimmer, 2020). Riparian/terrestrial wet spots are characterized by their positions at the freshwater–terrestrial transition zone or with high water availability/accumulation, encompassing a broad spectrum of habitats such as floodplains, riparian zones, and sites of groundwater seepage, among others (Selwood & Zimmer, 2020). Contrary to the spatially isolated climate relics (Woolbright et al., 2014) or insular ecosystems (Cartwright, 2019), river–ocean continuums (Wang et al., 2024) often give rise to vast climatic refugia due to their high connectivity, which can form cross-protection (Cloern et al., 2014; Gittman et al., 2016; Blanche et al., 2024).

Regardless of their vastness, a refugium cannot safeguard all types of life; indeed, surveys on diverse organisms might even yield contradictory evidence (Heusser, 1971). The majority of research concerning the biological conservation of refugia has concentrated on large animals, plants and invertebrates, with extremely scant attention given to microorganisms (Selwood & Zimmer, 2020). Microorganisms, too, can be sheltered by refugia (Mikryukov et al., 2021; Jackson et al., 2022). The extinction of all mammals would have little impact on the functioning of most ecosystems (Lawton, 2023), whereas the microbial extinctions would provoke unparalleled repercussions (Averill et al., 2022; Luo et al., 2024). Microbes can also be extinct (Averill et al., 2022), and the extinction rate is not slow (Louca et al., 2018). However, it is virtually impossible to accurately gauge even the local microbial extinction in a given refugium. So, determining how well a given refugium is persisting and an ecosystem is functioning requires quantifying the dynamics of species identity within the microbial community by leveraging suitable methodologies (Tokeshi, 1990).

Regarding the same refugium across different time points, the microbial community composition may be quite dissimilar (Vila et al., 2020). These “kaleidoscopic” changes in composition over annual timescales are attributable to the relatively temporary assemblages of species with distinct spatiotemporal dynamics in macroecology (Mac Nally et al., 2014). This impedes studying the dynamics of refugia, primarily due to the substantial prevalence of non-interacting rare species, which, in most cases, are the most vulnerable to extinction (Mac Nally, 2007). According to global relative abundance, only 0.4% of bacterial species can be classified as common worldwide (Bickel & Or, 2021). Upon ranking the entire microbial taxa in descending order of their abundance (i.e., the rank abundance curve, RAC), one will encounter a very steep curve accompanied by an elongated right tail (Mouquet & Loreau, 2003; McGill et al., 2007). The left side of the RAC corresponds to the more common species, while the right one corresponds to the rarer (Bickel & Or, 2021).

This pattern of species abundance distribution (SAD) engenders multi-faceted measurements of the internal dynamical processes in a community (Morlon et al., 2009; Avolio et al., 2019), helping to characterize the time trends inherent in refugia (Bennett & Provan, 2008; Avolio et al., 2015). Of these, the two most fundamental measures are changes in rank (the positioning of species abundance hierarchy) and in evenness (the steepness of the overall curve) (Collins et al., 2008). Rank reordering and the abundance of common species (an important regulator of evenness) are presumed to exert a more profound influence on community dynamics and ecosystem function than richness (Winfree et al., 2015; Jones et al., 2017). By further comparison, the effect of rank change is greater than that of evenness change (Wohlgemuth et al., 2016) and affects community stability (i.e., temporal Taylor’s slope) as well (Ghosh & Matthews, 2024). All these reports highlight the key role that rank hierarchy plays in determining refugial capacity, a role that is yet underappreciated in the context of microbial communities.

To this end, we investigate one-year monthly soil microbial community dynamics in three large-river deltaic estuaries (LDEs) in China, alongside examining a headwater and an inland riparian zone (Figure 1). Contrary to ordinary rivers, large rivers are widely recognized as climatic refugia, due to their complex lateral and longitudinal habitats and high anti-disturbance capabilities (Nilsson et al., 2005; Pyron et al., 2020). As one of the three mechanisms that foster resilience in river ecosystems, the refugia in large-river watersheds are ubiquitous (Van Looy et al., 2019). Conceptually, large rivers are classic systems of dendritic networks (other networks also exist; Benda et al., 2004), where energy and material flux ultimately gather in estuaries (Naiman & Décamps, 1997; Tonkin et al., 2018). Large-river estuaries represent the final node of a dendritic network characterized by a high number of nodes and branches, which promotes the maintenance of genetic diversity and indicates a relatively high potential for refugia formation (Morrissey et al., 2009; Henriques-Silva et al., 2019). Additionally, individuals in a dendritic river network can move and colonize between branches of the network (out-of-network; Campbell Grant et al., 2007). All of the above suggest that LDEs may be promising climatic refugia for microbial communities.

**Figure 1.**
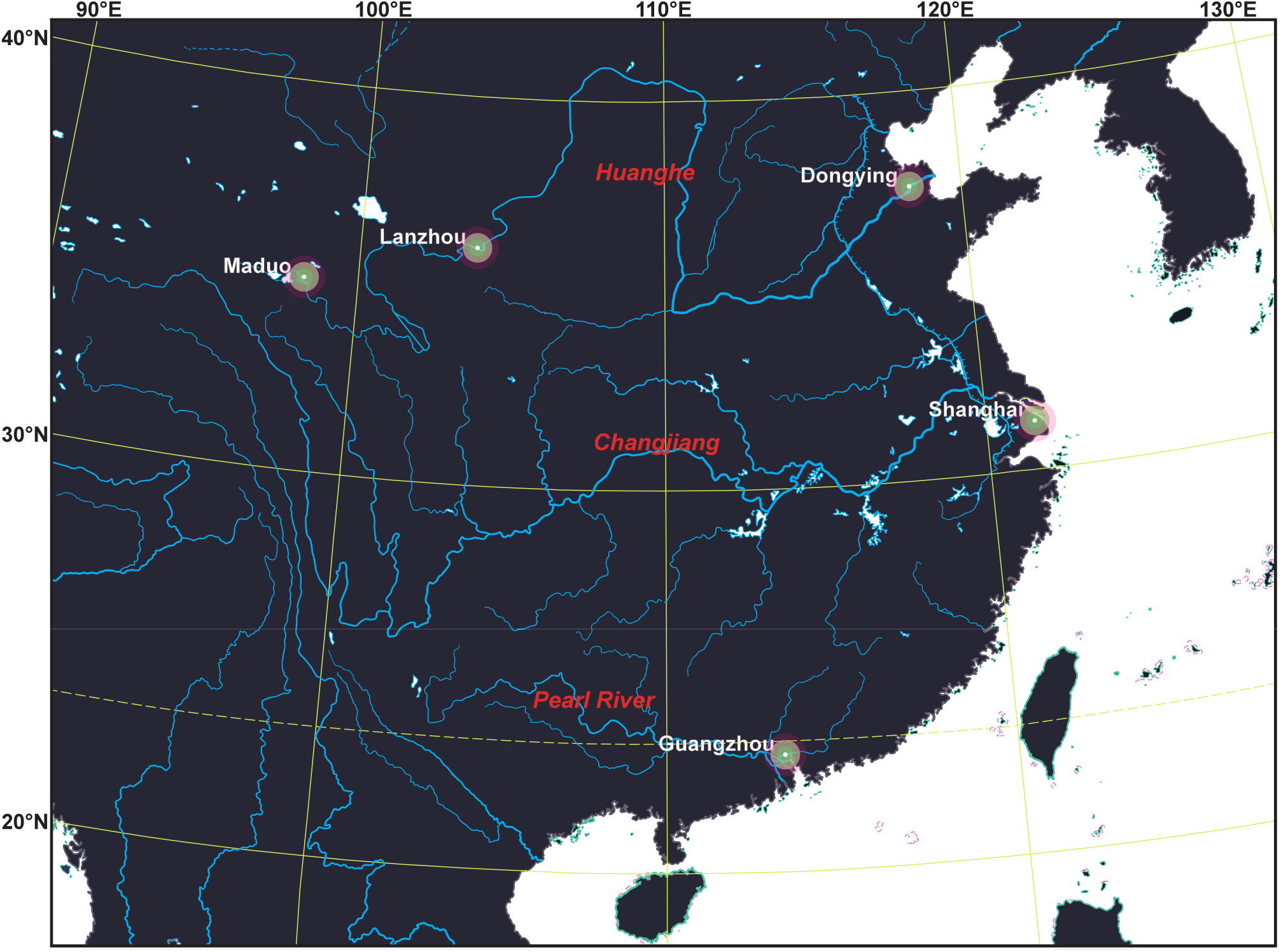
Sampling site map, located at the large-river deltaic estuaries (LDEs) of the first-order streams: Huanghe (Dongying), Changjiang (Shanghai), and Pearl River (Guangzhou), along with an inland riparian zone (Lanzhou) and a headwater (Maduo).

LDEs serve as crucial interfaces for the exchange of water and materials among large rivers, lands, and oceans (Cloern & Jassby, 2012; Sheng et al., 2024), and are highly susceptible to climate change (Bianchi & Allison, 2009; Chan et al., 2024). Therefore, this paper examines the climatic refugia for microorganisms in LDEs and explores the factors that contribute to their establishment, laying a foundation both for future LDE and microbial conservation efforts.

## Methods

### Site description and data collection

From 2012 to 2013, monthly soil samples were collected from the deltaic estuaries of the Huanghe (Dongying; sample size = 13), Changjiang (Shanghai; 12), and Pearl River (Guangzhou; 11)—the three largest rivers in China and among the top 20 in the world (Bianchi & Allison, 2009; Liu et al., 2014)—over the course of one year (Figure 1). A similar sampling protocol was also conducted in two sites located in the Three-River Source Region (Maduo; 12) and Loess Plateau (Lanzhou; 11). The climate across the five sites varies greatly, with mean annual temperature (MAT) varying from -1.6°C to 22.8°C, and mean annual precipitation (MAP) varying from 204.8 mm to 1,589.0 mm, as detailed in Zheng et al. (2018). Our investigation into climatic refugia for microbial communities excludes riverscour ecosystems (Estes et al., 2023) located too close to rivers (and coasts), where the dispersal and assembly of microorganisms are significantly influenced by water flow (McHergui et al., 2014; Ye et al., 2023). Despite being refugia for certain species, such as heliophytic species (Estes et al., 2023), these habitats are locally ephemeral (Corlett & Tomlinson, 2020). To streamline the search for refugia, our study focused on topsoil, where the heterogeneity of different substrates can be more effectively assimilated by overlying sediments (Corlett & Tomlinson, 2020). Here, the impacts of climatic and physicochemical properties are more pronounced compared to those in larger river and marine substrates (Bahram et al., 2018).

The selection of environmental indicators was based on several factors that exerted the most significant control over microbial communities in topsoil, as outlined in Bahram et al. (2018). Topsoil samples (5 cm depth) were collected from each site at each sampling time, totaling 59 samples. Surface litter was removed prior to mixing the samples with the surrounding soil. Subsequently, the samples were promptly placed on dry ice and stored in a laboratory freezer at -20°C for subsequent processing. pH was measured three times with a pH meter (Mettler Toledo, USA) after thawing the soil sample at room temperature, mixing it with ultra-pure water in a ratio of 1:2.5 (w/v, g/mL), shaking for 30 minutes, and centrifuging for 10 minutes at 3000 r/min, with the mean value calculated. Total carbon and total nitrogen were quantified using an organic element analyzer EA1110 (Carlo Erba, Italy), following the method described by Zhang et al. (2012). The monthly mean temperature (MMT) and the monthly mean precipitation (MMP) were obtained in the earlier work using a variety of data collection methods (zheng et al., 2016).

To extract DNA from each sample, the FastDNA Spin Kit for Soil (MP Biomedicals, USA) was used according to the manufacturer’s protocol. Primers 338F (5′-ACTCCTACGGGAGGCAGCA-3′) and 806R (5′-GGACTACNVGGGTWTCTAAT-3′) were used to amplify the hypervariable region V3–V4 of bacterial 16S rRNA gene. Each polymerase chain reaction (PCR) amplification was conducted in a 20 μl reaction system using TransGen AP221-02: TransStart Fastpfu DNA Polymerase (TransGen Biotech, China). The reaction mixtures included 4 μl of 5× FastPfu Buffer, 2 μl of 2.5 mM dNTPs, 0.8 μl each of Forward Primer (5 μM) and Reverse Primer (5 μM), 0.4 μl of FastPfu Polymerase, 0.2 μl of BSA, 10 ng of Template DNA, and double distilled water added to make up the final volume to 20 μl. PCR amplification was performed using the ABI GeneAmp® 9700 PCR System (Applied Biosystems, USA) with the following protocol: an initial pre-denaturation step at 95°C for 3 min; followed by 27 cycles of denaturation at 95°C for 30 s, annealing at 55°C for 30 s, and extension at 72°C for 45 s; and a final extension step at 72°C for 10 min. We observed lower quality scores and a reduced sample size for paired-end sequencing from the reverse reads (3′ end, R2) generated by the Illumina MiSeq platform at Shanghai Majorbio Bio-Pharm Technology Co., Ltd. To resolve this issue, we chose single-end (forward reads, R1) sequencing, which effectively ameliorates the problems associated with low-quality reverse reads and yields results comparable to those of paired-end sequencing (Werner et al., 2012; Ramakodi, 2021). The DADA2 (Callahan et al., 2016) plugin in the QIIME2 (Bolyen et al., 2019) pipeline was used to denoise the sequencing reads and infer amplicon sequence variants (ASVs). Taxonomy assignment was performed using the qiime2 classify-sklearn plugin and the SILVA 132 database (Quast et al., 2013). For diversity calculations, the rarefaction depth was set at a minimum sequencing depth of 19,923 ASVs. The raw sequence data reported in this paper have been deposited in the Genome Sequence Archive (Chen et al., 2021) in National Genomics Data Center (CNCB-NGDC Members and Partners, 2023), China National Center for Bioinformation/Beijing Institute of Genomics, Chinese Academy of Sciences (GSA: CRA018508) that are publicly accessible at https://ngdc.cncb.ac.cn/gsa.

### Statistical analyses

Microbial samples exhibit greater biodiversity (i.e., a single large sample; McGill, 2003) compared to macroorganisms, resulting in SADs that are dominated by abiological mechanisms, such as statistical processes (McGill, 2003; Šizling et al., 2009). This can lead to an excessively steep RAC (Nemergut et al., 2013), which may obscure the dominance (rank) hierarchy and internal dynamics of the microbial community. As a typical composite community, co-occurrence within the microbial community may not necessarily imply complete interaction (Fauth et al., 1996; Nemergut et al., 2013). Instead, it more likely indicates the presence of multiple subcommunities with limited or zero net effect interactions and significant differences in assembly mechanisms (Fan et al., 2024; Srinivasan et al., 2024). This precisely means that composite communities are heterogeneous and should be subdivided to break up overly steep curves (MacArthur, 1957). Therefore, we employed the logistic-tree normal latent Dirichlet allocation (LTN-LDA) model (LeBlanc & Ma, 2023) to divide microbial subcommunities (or sub-assemblages; Hosoda et al., 2020) at the genus level, aiming to identify the dominant and rare genera within each subcommunity. This model simplifies the taxonomic complexity and enables the investigation of microbial ecologically driven RAC shape dynamics at a lower level of biodiversity and a more localized scale (Sankaran & Holmes, 2019).

As a mixed-membership model, the LDA was initially developed for topic discovery in natural language processing (Blei et al., 2003; Griffiths & Steyvers 2004) and has since been used in microbial ecology to identify taxa that co-occur and exhibit consistent dynamic changes, as well as to find subcommunities (Shafiei et al., 2015; Higashi et al., 2018). Unlike traditional clustering methods, the LDA model assigns a mixture probability to each subcommunity, making it more compatible with real-world scenarios. For instance, a species can be dominant in multiple months or across various plots, rather than being confined to just one. Let the *W* genera from all *D* samples be divided into *K* subcommunities. To obtain any genus *w*_*dn*_, the following steps are taken: First, a subcommunity distribution, *θ*_*d*_, for the *d* th sample is sampled from the Dirichlet distribution *θ*_*d*_ *∼* Dirichlet *α* . Next, a subcommunity *z*_*dn*_ is obtained from the multinomial distribution *z*_*dn*_ | *θ*_*d*_ *∼* Multinomial *θ*_*d*_ to determine the subcommunity to which the genus *w*_*dn*_ belongs. Subsequently, the genus distribution *β*_*k*_ for subcommunity *k* is sampled from the Dirichlet distribution *β*_*k*_ *∼* Dirichlet *γ* . Finally, this genus is obtained from the multinomial distribution 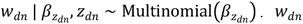. *w*_*dn*_ is the genus corresponding to the *n* th sequencing read in the *d* th sample (Hosoda et al., 2020; Deek & Li, 2021). *α* and *γ* represent the hyperparameters for the sample–subcommunity and subcommunity–genus Dirichlet distributions, respectively.

To solve the problem of significant heterogeneity among different samples, logistic tree normal (LTN) was integrated into the LDA model, leveraging the inherent phylogenetic tree of microbial communities (Wang et al., 2022; LeBlanc & Ma, 2023). It was assumed that taxa closer to the leaf nodes of the phylogenetic tree exhibit greater heterogeneity across samples compared to those further away. The rationale behind this assumption is that taxa close to each other at deep levels of the phylogenetic tree tend to exhibit greater functional similarity and competitiveness (Tang et al., 2018; Jeganathan & Holmes, 2021; LeBlanc & Ma, 2023). The tuning parameter *C* was used to divide the phylogenetic tree into upper and lower parts, with the lower part suggesting significant cross-sample heterogeneity. Using the LTNLDA package in R (LeBlanc & Ma, 2023), we initially excluded taxa that appeared fewer than 100 times across all samples. Then we performed 1,000 Gibbs sampling iterations, with a burn-in period of 10,000, for various pairs of *K* and *C* within the appropriate range, assessing performance using perplexity (Wallach et al., 2009). Finally, we used 5 × 2-fold cross-validation to determine the optimal values for *K* and *C* (Dietterich, 1998), resulting in *K* = 5 and *C* = 9 (Appendix S1: Table S1).

The RAC at each sampling time for each site was composed of five selected subcommunities (*K*), and the evenness index *E*_*var*_ proposed by Smith & Wilson (1996):

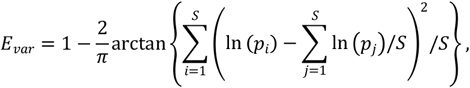

Where *S* is the number of subcommunities in a given sample, and *p*_*i*_ is the relative abundance of the *i*th subcommunity (the sum of all genera within). The evenness change Δ*E* is the difference of *E*_*var*_ between the sampling times before and after (Avolio et al., 2019). The rank change or mean rank shift Δ*R* is calculated as: 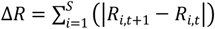. Where *R*_*i*,*t*_ is the rank of species *i* at time *t*, and *R*_*i*,*t* 1_ is the rank at *t* +1 (Collins et al., 2008).

The sampling dates in this study were not on the same day of every month and differed among sites in terms of both the starting and ending months (i.e., varying time intervals; Hasl et al., 2024). To remedy this issue, We used a hierarchical Bayesian continuous time dynamic model, which combines a hierarchical random-effects model with a continuous time dynamic model (Driver & Voelkle, 2018). This approach enables us to convert irregular time intervals from bias into valuable sampling replicates, thereby increasing the sample size and enhancing the predictive accuracy of time dynamics (Hasl et al., 2024). In simple terms, the model eliminates time-interval dependency by treating time as continuous rather than discrete and uses the Bayesian hierarchical approach to share the structure of the model among samples across all time and sites, so that variables from all other samples serve as priors for parameter estimation of variables for each sample (i.e., full random effects; Driver & Voelkle, 2018; Driver & Tomasik, 2023). The stochastic differential equation applied in this paper is (Lins Machado & He, 2024): d***y*** (*t*) = (**A*y*** (*t*) **b**)d*t* **G**d**W** (*t*) . The vector ***y*** *t* is the state of all latent processes at time *t* and **b** is a continuous time intercept. The drift matrix **A** exhibits auto effects on the diagonal elements and cross effects on the off-diagonals. d**W** (*t*) is the stochastic noise and **G** represents its effect. Auto effects refer to how each variable influences its own future value, whereas cross effects denote the influence of one variable on the future value of another. Each variable was grand mean centered and scaled (Driver et al., 2017), and each manifest and latent variable had the same name (Lins Machado & He, 2024). The hierarchical Bayesian continuous time dynamic model was implemented through the Bayesian extension of the ctsem package in R (Driver et al., 2017).

## Results

The five subcommunities comprised a total of 477 genera, and the five genera with the highest relative abundance in each subcommunity were displayed (Figure 2). The minimum number of genera required to account for half of the total relative abundance in each subcommunity was as follows: subcommunity 1, 13 (2.7%); subcommunity 2, 11 (2.3%); subcommunity 3, 13 (2.7%); subcommunity 4, 2 (0.4%); and subcommunity 5, 1 (0.2%). The five subcommunities are ranked from 1 to 5 based on the mean relative abundance of all 59 samples across the five sites, reflecting the overall status of the five subcommunities (Figure 3). However, the ranking might differ in certain samples. Subcommunities 1 and 2 were present in all sites, while subcommunities 3 and 4 were only visibly present in Dongying and Guangzhou, respectively (Figure 3). 91% of subcommunity 5 is composed of the single genus *Massilia* and was only evident in some samples from Shanghai (Figures 2 and 3).

**Figure 2.**
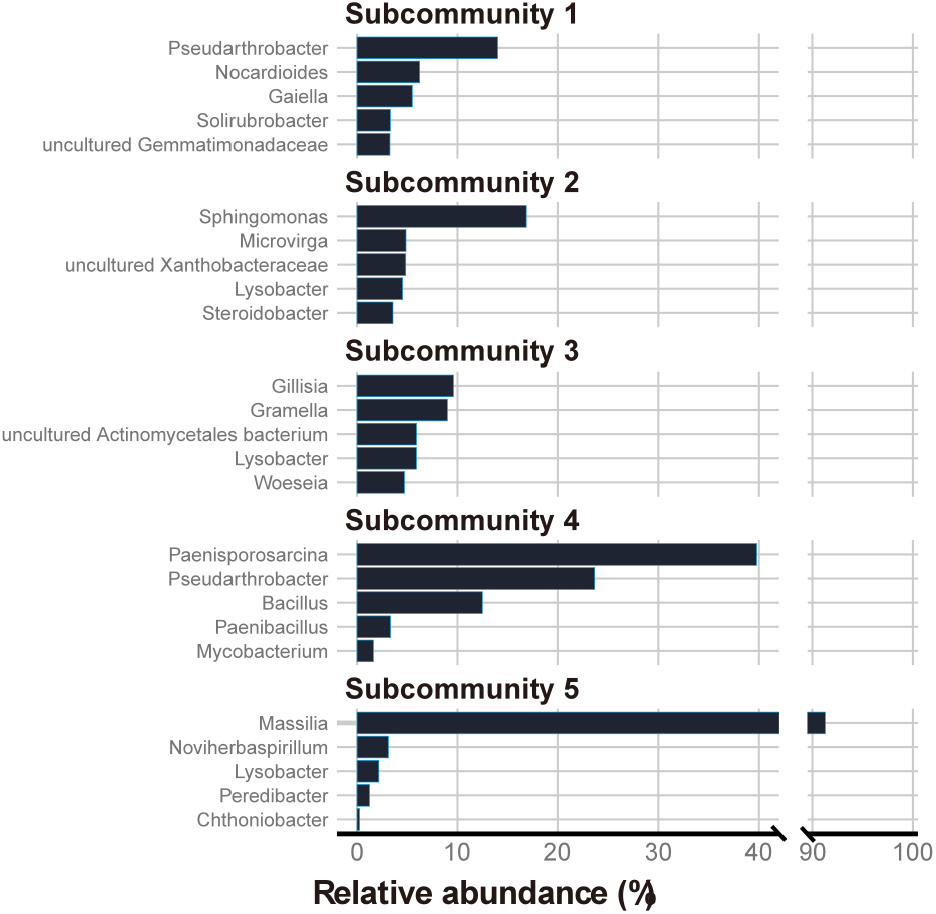
The top five genera with the highest relative abundance in five subcommunities. The X-axis is truncated to accommodate the relative abundance of genus *Massilia*.

**Figure 3.**
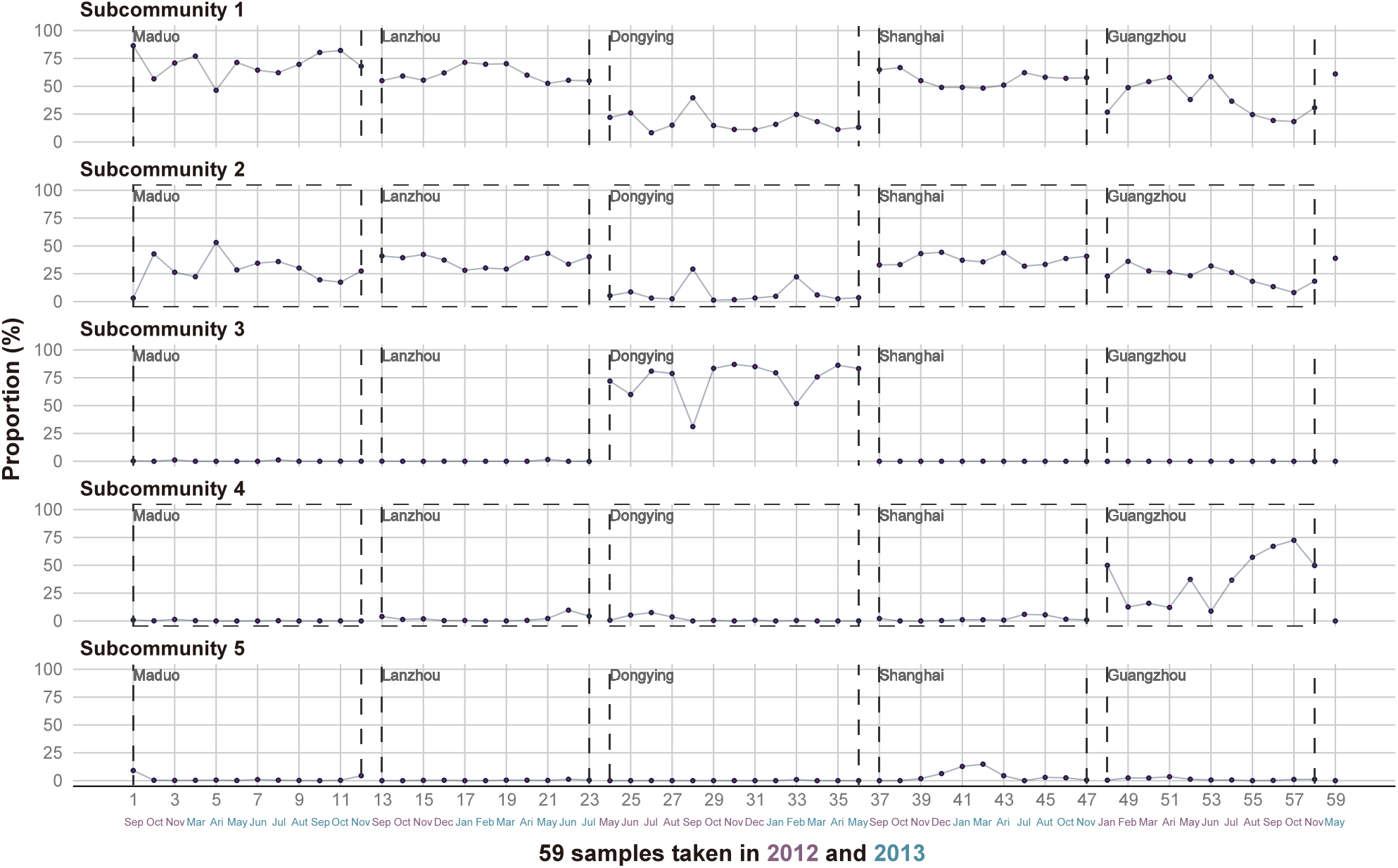
The distribution proportions of five subcommunities across 59 different sampling times at five sites, with the sum of the proportions of the five subcommunities in each column equal to one. All samples were collected in 2012 and 2013, with the sampling months and years displayed at the bottom. Each box composed of long-dashed lines belongs to a sampling site, and the sample numbers connected to adjacent boxes are not related. The 59th sample belongs to Shanghai; it was not changed to maintain consistency with the original data numbering.

The RAC and alpha diversity for each sample were displayed (Figure 4). For alpha diversity, although individual samples may show higher or lower values, there are no significant differences overall among different sites. For RAC, first, there were relatively consistent rank hierarchy differences between different sites. Secondly, results from different sampling times within the same site were inconsistent. However, the variations in RACs seemed to revolve around a constant hierarchical pattern. Therefore, although there were usually some differences in rank or evenness between adjacent samples’ RACs, the RACs from the earliest and latest sampling times for each site remained almost unchanged.

**Figure 4.**
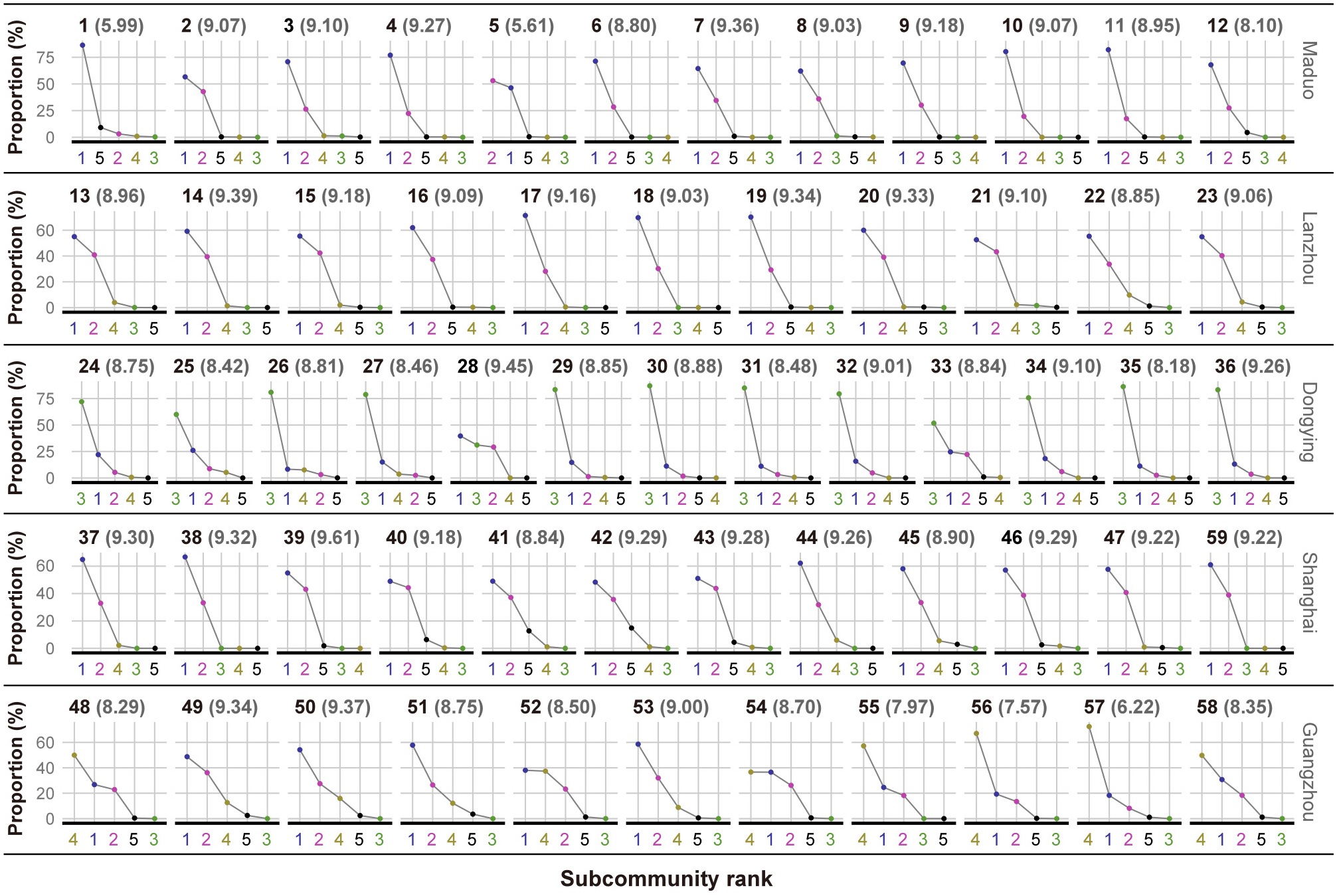
Figure 4 The rank abundance curves (RACs) of all samples for the subcommunities, with numbers on the abscissa from 1 to 5, corresponding to subcommunities 1 to 5. The numbers in parentheses after each sample number represent Shannon entropy (i.e., alpha diversity).

Based on the predicted trajectories of all observations (Figure 5; Appendix S1: Table S2), Δ***R*** and Δ***E*** fluctuated around the almost same baseline across different sites, while the predicted trajectories of MMT, MMP, soil carbon to nitrogen ratio (C/N), and pH differed among the various sites. Among them, Maduo and Lanzhou represented the maximum or minimum predicted values for the variables, while Guangzhou represented significantly opposite predicted values. The predicted values for Dongying and Shanghai were situated in the moderate range. From the pH, the clearest separation of LDEs was observed. From posterior intervals and point estimates (50% median) for means of population distribution estimated by the hierarchical Bayesian continuous time dynamic model, it was observed that, except for the auto effect parameters of the drift matrix, other parameters did not show significant differences. Qualitatively similar results were obtained when only LDE sites were considered and latitude was set as a time-independent predictor. The significant negative value of the automatic effect is pervasive, indicating that the variable’s negative feedback mechanism maintains stability over time. The point estimates for the auto effect parameters, sorted from largest to smallest, were as follows: MMT, -1.6543; MMP, -2.1549; Δ***R***, -2.2519; pH, -2.7709; Δ***E***, -3.2408; and C/N, -3.528. The smaller (i.e., closer to zero) the negative value of a variable, the less the negative feedback pressure to correct it back to the expected baseline in the next moment when it deviates from the anticipated trend. This means that a single change in the variable can have a longer-lasting impact on itself. Conversely, the larger the negative value of a variable, the stronger the pressure to correct it back to the trend, meaning that a single change in the variable (such as a one-time disturbance) can only cause a short-term effect. For our results, the minimum negative value of MMT indicated that its single change caused the most lasting consequences, while C/N was the opposite (Appendix S2).

**Figure 5.**
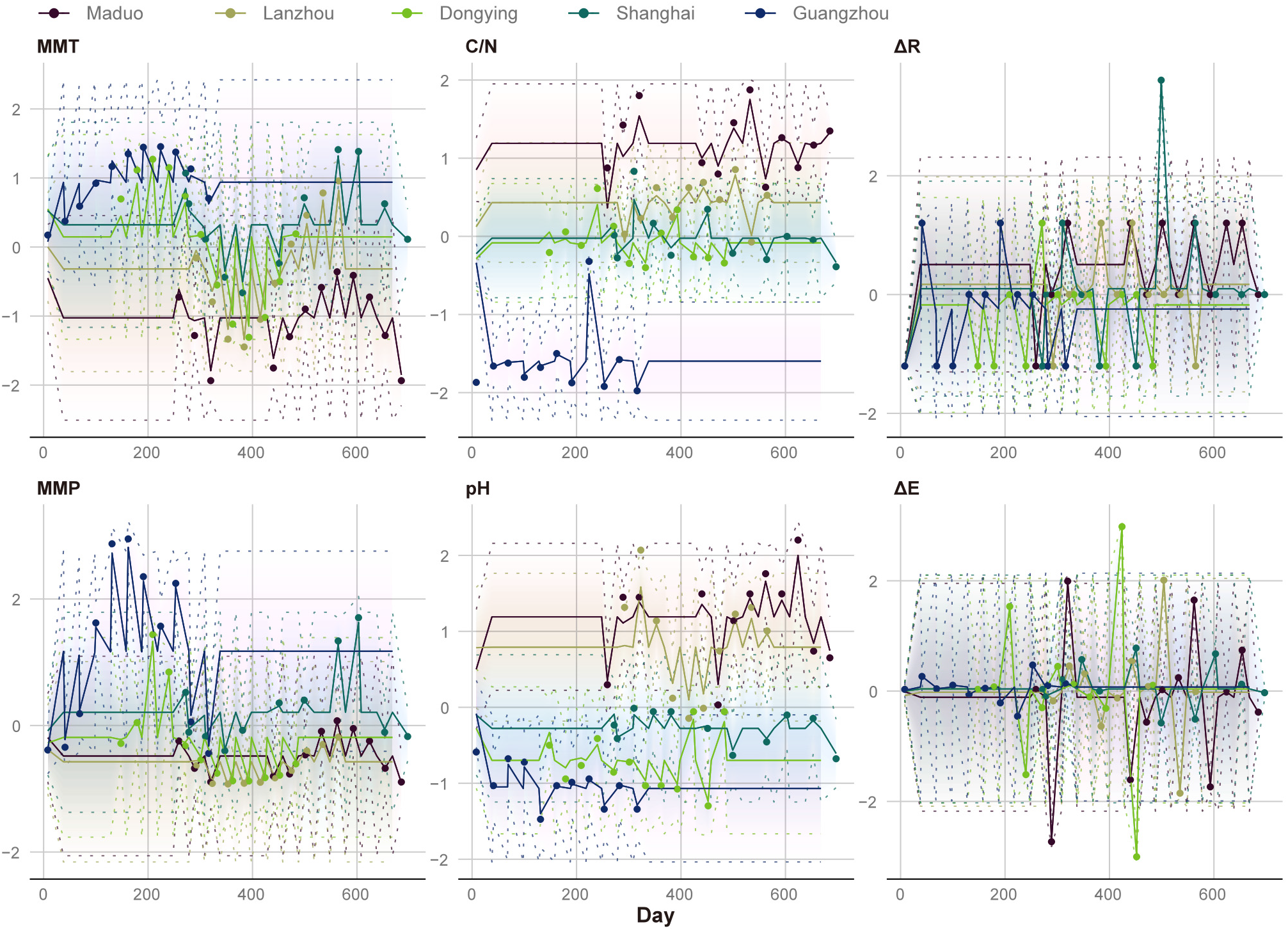
The observed scores (points) of different variables at different sampling times and the consequently predicted scores (solid lines) from the Kalman smoother, which reduce errors caused by uncertainty. The 95% credible interval (dashed lines) represents the uncertainty. The abscissa of the observed scores represents sampling times accurate to the day. Observations start from the earliest sampling date of the Guangzhou sample on January 8, 2012 (day 8) and continue until the latest sampling date of the Shanghai sample on November 27, 2013 (day 697). Abbreviation: MMT, monthly mean temperature; MMP, monthly mean precipitation; C/N, soil carbon to nitrogen ratio; pH, soil pH; Δ***R***, rank change; Δ***E***, evenness change.

## Discussion

We have found that LDEs act as apparent climatic refugia for microbial communities, with each LDE exhibiting a distinct rank hierarchy of subcommunities influenced by climate and soil conditions, and maintaining constancy over time. Although we know little about the rules governing extinction and survival (Jablonski, 2005), the results from this study, viewed from the perspective of microbial subcommunities, are highly consistent with ecological processes observed in macroorganisms. This implies that numerous potential strategies could be applied to protect microbes.

Each LDE is unique, yet they still have commonness. We found that the same subcommunities (1 and 2) appear throughout the large-river watersheds and in different LDEs, although their proportion may be lower in some LDEs. This background soil microbial subcommunity, known as the core floodplain microbiome, which persists across the entire watershed on an annual timescale, has been confirmed and explained by recurring scaling motifs in river networks from the micrometer to the watershed scale (Matheus Carnevali et al., 2021). Additionally, these background subcommunities control substantial changes in niches within river networks and have similar ecological requirements, which may be a reason for their homogenous selection (Hui et al., 2023). For subcommunities that are rich only in LDEs (3–5), it seems that only a case-by-case analysis can be performed. Although our results showed stable microbial hierarchies, taxa that adapted differently to climate—even in contrasting ways—might have periodically retreated to refugia during cyclical climate changes (Stewart et al., 2010). They may switch between LDE-affiliated and background species, and species identities may also shift between the umbrella species and the protected ones (Andelman & Fagan, 2000). Finally, whether LDEs are widely occupied by soil microorganisms due to their favorable refuge conditions, or if they are merely a complex of widely distributed, isolated microrefugia, remains undetermined (Rull, 2009, 2010). Therefore, the geographically uneven distribution of biodiversity may be intrinsic rather than a result of external influences (Rauch & Bar-Yam, 2004).

From the perspective within subcommunities, although the composite communities have been partitioned, our results still show that half of the relative abundance of each subcommunity is occupied by a very small number of genera (0.2%–2.7%). This phenomenon, known as hyperdominance (ter Steege et al., 2014), illustrates the huge gap between common and rare species and is reflected in the RACs. The strongly right-skewed shape of the RAC is believed to be caused by non-ecological mechanisms, such as the central limit theorem (Šizling et al., 2009), because no factors that could significantly affect the shape of the RAC have been identified (Nekola & Brown, 2007). For example, a large-scale study collecting 18 environmental variables known to have significant effects on bird species found that none were associated with the shape of the RAC for bird assemblages (Yen et al., 2013). Similarly, our results also confirmed that the four variables considered to have the greatest impact on shaping global topsoil microbial communities (MMT, MMP, C/N, and pH; originally MAT and MAP) were unrelated to changes in the shape of the RAC. Although curves shaped under different diversity conditions might offer varying observational windows for RACs (Locey & White, 2013), it is nearly impossible for SADs at similar taxonomic levels in microbes to be completely unveiled, like those of the Insecta (Callaghan et al., 2023). In addition, changes in biodiversity do not necessarily lead to changes in rank (Dornelas et al., 2014).

In fluctuating environments, extinction events of microbes are somewhat repeatable (Rodríguez-Verdugo & Ackermann, 2020). Although local extinctions of rare species within different subcommunities are likely still random, resembling a “field of bullets,” the impact on the loss of evolutionary history is not significant (Nee & May, 1997; Faller et al., 2008). Therefore, the dominance structure of specific subcommunities in LDE refugia might also be a deterministic outcome, maintaining ecological inertia through long-term climatic fluctuations (Stralberg et al., 2020). In this study, we showed that changes in rank structure had an impact on themselves that was second only to climate, while changes in evenness had a shorter-term impact on themselves. Although the reordering of ranks among rare subcommunities (on the right side of RACs) should theoretically be frequent, our observational evidence suggests otherwise. This self-stabilizing negative feedback might not be detectable from the correlational method alone and is not necessarily confined to climatic refugia. In deed, some rare species with high local extinction risks carry high functional vulnerability, but they support unique ecosystem functions, which confirms the ecological significance of changes in evenness (Petchey et al., 2008; Mouillot et al., 2013; Leitão et al., 2016). However, the microbial community dynamics caused by rank changes, comparable to community succession, may need to be studied on a timescale of a century (Magurran, 2007; Dini-Andreote et al., 2014).

Due to the large scale of microbial SAD, the high diversity, and the low niche dimensionality, understanding their fine-scale ecological processes may be redundant (Chisholm & Pacala, 2010). Larger microbial communities can be viewed as composed of smaller neutral subcommunities for each niche (van Nes et al., 2024), which helps to generalize microbial ecology and supports the conservation of climatic refugia. The stable existence of microorganisms is also one of the foundations for the survival of other organisms in their refugia.

## Acknowledgments

This work was supported by the Project of Southern Marine Science and Engineering Guangdong Laboratory (Zhuhai) (No. SML2023SP218), the National Natural Science Foundation of China (grant Nos. 92051117, 41776137), the National Key Basic Research Program of China (grant No. 2022YFC2805505), and the Guangdong Basic and Applied Basic Research Foundation (grant No. 2021B1515120080). We acknowledge the contributions of Yingqin Wu, Bangqi Hu, and Joanna Zhang for their assistance with field sampling, as well as Yarong Feng and Carolina Lins Machado for their valuable comments on the manuscript.

## Data Availability Statement

The data that support the findings of this study are openly available in National Genomics Data Center at https://ngdc.cncb.ac.cn/gsa (GSA: CRA018508).

