## Appendix S2 for "Microbes under climatic refugia: Self-stabilizing subcommunity rank dynamics in large-river deltaic estuaries"

Subject parameter distribution:

$$\underbrace{\begin{bmatrix} \text{V1_MMT_t0}_i \\ \text{V2_CN_t0}_i \\ \text{V3_Rank_t0}_i \\ \text{V4_MMP_t0}_i \\ \text{V5_pH_t0}_i \\ \text{V6_Evenness_t0}_i \\ \text{mm_V1_MMT}_i \\ \text{mm_V2_CN}_i \\ \text{mm_V3_Rank}_i \\ \text{mm_V4_MMP}_i \\ \text{mm_V5_pH}_i \\ \text{mm_V6_Evenness}_i \\ \text{V1_MMT_cint}_i \\ \text{V2_CN_cint}_i \\ \text{V3_Rank_cint}_i \\ \text{V4_MMP_cint}_i \\ \text{V5_pH_cint}_i \\ \text{V6_Evenness_cint}_i \end{bmatrix}}_{\boldsymbol{\phi}(i)} \approx \mathrm{N}\left( \begin{bmatrix} 0.171 \\ -0.253 \\ -1.453 \\ -0.089 \\ 0.14 \\ -0.038 \\ 0.038 \\ 0.021 \\ 0.25 \\ -0.112 \\ -0.086 \\ 0.029 \\ -0.147 \\ -0.062 \\ -0.352 \\ 0.378 \\ 0.265 \\ -0.069 \end{bmatrix}, \begin{bmatrix}
0.231 & 0.015 & 0 & 0.05 & -0.018 & 0.002 & -0.006 & -0.016 & 0 & 0.014 & -0.048 & 0.003 & 0.116 & -0.06 & -0.057 & -1.552 & -0.343 & 0.026 \\
0.015 & 0.053 & 0 & 0.011 & -0.002 & 0.001 & 0.007 & 0.006 & 0 & 0.011 & -0.004 & 0.002 & -0.061 & 0.038 & 0.016 & -1.042 & 0.069 & -0.004 \\
0 & 0 & 0.001 & 0 & 0 & 0 & 0 & 0 & 0 & 0 & 0 & 0 & 0 & 0 & 0 & -0.004 & -0.002 & 0 \\
0.05 & 0.011 & 0 & 0.148 & -0.004 & 0.001 & 0.009 & 0.029 & 0 & -0.02 & -0.01 & 0.002 & -0.022 & -0.029 & 0.034 & -0.072 & 0.061 & 0.012 \\
-0.018 & -0.002 & 0 & -0.004 & 0.229 & -0.007 & -0.001 & 0.04 & 0 & -0.017 & -0.005 & -0.006 & 0.017 & 0.01 & -0.016 & -0.823 & 0.296 & 0.022 \\
0.002 & 0.001 & 0 & 0.001 & -0.007 & 0.008 & 0.001 & -0.001 & 0 & 0.004 & -0.011 & -0.002 & -0.003 & -0.005 & 0.001 & 0.094 & 0.005 & 0.003 \\
-0.006 & 0.007 & 0 & 0.009 & -0.001 & 0.001 & 0.275 & -0.089 & 0 & 0.047 & -0.076 & 0.001 & -0.068 & -0.03 & -0.043 & 2.918 & -0.351 & 0.029 \\
-0.016 & 0.006 & 0 & 0.029 & 0.04 & -0.001 & -0.089 & 0.534 & 0 & -0.01 & 0.139 & -0.005 & -0.117 & -0.031 & 0.122 & -4.405 & 1.036 & -0.028 \\
0 & 0 & 0 & 0 & 0 & 0 & 0 & 0 & 0.001 & 0 & 0 & 0 & 0 & 0 & 0 & -0.001 & 0 & 0 \\
0.014 & 0.011 & 0 & -0.02 & -0.017 & 0.004 & 0.047 & -0.01 & 0 & 0.156 & -0.035 & 0.003 & 0.038 & -0.022 & 0.01 & -0.495 & -0.021 & -0.003 \\
-0.048 & -0.004 & 0 & -0.01 & -0.005 & -0.011 & -0.076 & 0.139 & 0 & -0.035 & 0.352 & -0.01 & -0.068 & 0.06 & 0.113 & -2.685 & 0.118 & -0.019 \\
0.003 & 0.002 & 0 & 0.002 & -0.006 & -0.002 & 0.001 & -0.005 & 0 & 0.003 & -0.01 & 0.009 & -0.001 & -0.001 & -0.002 & 0.118 & -0.001 & -0.005 \\
0.116 & -0.061 & 0 & -0.022 & 0.017 & -0.003 & -0.068 & -0.117 & 0 & 0.038 & -0.068 & -0.001 & 6.85 & -0.046 & -0.092 & 17.606 & -0.275 & 0.005 \\
-0.06 & 0.038 & 0 & -0.029 & 0.01 & -0.005 & -0.03 & -0.031 & 0 & -0.022 & 0.06 & -0.001 & -0.046 & 1.022 & 0.071 & -0.14 & 0.147 & -0.033 \\
-0.057 & 0.016 & 0 & 0.034 & -0.016 & 0.001 & -0.043 & 0.122 & 0 & 0.01 & 0.113 & -0.002 & -0.092 & 0.071 & 2.653 & -1.001 & 0.491 & -0.021 \\
-1.552 & -1.042 & -0.004 & -0.072 & -0.823 & 0.094 & 2.918 & -4.405 & -0.001 & -0.495 & -2.685 & 0.118 & 17.606 & -0.14 & -1.001 & 718.297 & -4.466 & 0.792 \\
-0.343 & 0.069 & -0.002 & 0.061 & 0.296 & 0.005 & -0.351 & 1.036 & 0 & -0.021 & 0.118 & -0.001 & -0.275 & 0.147 & 0.491 & -4.466 & 25.832 & -0.136 \\
0.026 & -0.004 & 0 & 0.012 & 0.022 & 0.003 & 0.029 & -0.028 & 0 & -0.003 & -0.019 & -0.005 & 0.005 & -0.033 & -0.021 & 0.792 & -0.136 & 1.995
\end{bmatrix} \right)$$

Initial latent state:

$$\underbrace{\begin{bmatrix} \text{V1\_MMT} \\ \text{V2\_CN} \\ \text{V3\_Rank} \\ \text{V4\_MMP} \\ \text{V5\_pH} \\ \text{V6\_Evenness} \end{bmatrix}(t_0)}_{\boldsymbol{\eta}(t_0)} \sim \mathrm{N}\left( \underbrace{\begin{bmatrix} 0.171 \\ -0.253 \\ -1.453 \\ -0.089 \\ 0.14 \\ -0.038 \end{bmatrix}}_{\substack{\\ \text{T0MEANS}}}, \underbrace{UcorSDtoCov\left\{ \begin{bmatrix} 0.041 & 0 & 0 & 0 & 0 & 0 \\ 0.184 & 0.018 & 0 & 0 & 0 & 0 \\ 0.019 & 0.021 & 0.001 & 0 & 0 & 0 \\ 0.285 & 0.119 & 0.006 & 0.032 & 0 & 0 \\ -0.071 & -0.019 & -0.014 & -0.015 & 0.038 & 0 \\ 0.05 & 0.048 & 0.004 & 0.015 & -0.211 & 0.008 \end{bmatrix} \right\}}_{\substack{\mathbf{Q^*}_{t0} \\ \text{T0VAR}}} \right)$$

Deterministic change:

$$\underbrace{\mathrm{d}\begin{bmatrix} \text{V1\_MMT} \\ \text{V2\_CN} \\ \text{V3\_Rank} \\ \text{V4\_MMP} \\ \text{V5\_pH} \\ \text{V6\_Evenness} \end{bmatrix}(t)}_{\mathrm{d}\boldsymbol{\eta}(t)} = \left( \underbrace{\begin{bmatrix} -2.473 & -0.042 & -0.184 & 0.693 & -0.415 & 0.033 \\ -0.09 & -3.854 & 0.01 & -0.1 & 0.159 & -0.042 \\ -0.199 & 0.065 & -2.723 & -0.11 & 0.614 & -0.061 \\ 0.073 & 0 & -0.039 & -7.462 & -0.326 & 0.026 \\ -0.161 & 0.041 & 0.037 & -0.255 & -3.958 & -0.088 \\ 0.074 & -0.037 & -0.034 & -0.03 & -0.192 & -3.827 \end{bmatrix}}_{\substack{\mathbf{A} \\ \text{DRIFT}}} \underbrace{\begin{bmatrix} \text{V1\_MMT} \\ \text{V2\_CN} \\ \text{V3\_Rank} \\ \text{V4\_MMP} \\ \text{V5\_pH} \\ \text{V6\_Evenness} \end{bmatrix}(t)}_{\boldsymbol{\eta}(t)} + \underbrace{\begin{bmatrix} -0.147 \\ -0.062 \\ -0.352 \\ 0.378 \\ 0.265 \\ -0.069 \end{bmatrix}}_{\substack{\mathbf{b} \\ \text{CINT}}} \right) \mathrm{d}t \quad +$$

Random change:

$$\underbrace{UcorSDtoChol\left\{ \begin{bmatrix} 1.785 & 0 & 0 & 0 & 0 & 0 \\ 0.036 & 1.036 & 0 & 0 & 0 & 0 \\ 0.11 & -0.085 & 2.159 & 0 & 0 & 0 \\ 0.333 & 0.088 & 0.083 & 5.952 & 0 & 0 \\ 0.353 & -0.006 & -0.217 & -0.008 & 1.381 & 0 \\ -0.067 & 0.054 & 0.061 & 0.053 & 0.29 & 2.981 \end{bmatrix} \right\}}_{\substack{\mathbf{G} \\ \text{DIFFUSION}}} \underbrace{\mathrm{d}\begin{bmatrix} W_1 \\ W_2 \\ W_3 \\ W_4 \\ W_5 \\ W_6 \end{bmatrix}(t)}_{\mathrm{d}\mathbf{W}(t)}$$

Observations:

$$\underbrace{\begin{bmatrix} \text{V1\_MMT} \\ \text{V2\_CN} \\ \text{V3\_Rank} \\ \text{V4\_MMP} \\ \text{V5\_pH} \\ \text{V6\_Evenness} \end{bmatrix}(t)}_{\mathbf{Y}(t)} = \underbrace{\begin{bmatrix} 1 & 0 & 0 & 0 & 0 & 0 \\ 0 & 1 & 0 & 0 & 0 & 0 \\ 0 & 0 & 1 & 0 & 0 & 0 \\ 0 & 0 & 0 & 1 & 0 & 0 \\ 0 & 0 & 0 & 0 & 1 & 0 \\ 0 & 0 & 0 & 0 & 0 & 1 \end{bmatrix}}_{\substack{\boldsymbol{\Lambda} \\ \text{LAMBDA}}} \underbrace{\begin{bmatrix} \text{V1\_MMT} \\ \text{V2\_CN} \\ \text{V3\_Rank} \\ \text{V4\_MMP} \\ \text{V5\_pH} \\ \text{V6\_Evenness} \end{bmatrix}(t)}_{\boldsymbol{\eta}(t)} + \underbrace{\begin{bmatrix} 0.038 \\ 0.021 \\ 0.25 \\ -0.112 \\ -0.086 \\ 0.029 \end{bmatrix}}_{\substack{\boldsymbol{\tau} \\ \text{MANIFESTMEANS}}} \quad +$$

Observation noise:

$$\underbrace{\begin{bmatrix} 0.271 & 0 & 0 & 0 & 0 & 0 \\ 0 & 0.123 & 0 & 0 & 0 & 0 \\ 0 & 0 & 0.009 & 0 & 0 & 0 \\ 0 & 0 & 0 & 0.202 & 0 & 0 \\ 0 & 0 & 0 & 0 & 0.209 & 0 \\ 0 & 0 & 0 & 0 & 0 & 0.045 \end{bmatrix}}_{\substack{\boldsymbol{\Theta} \\ \text{MANIFESTVAR}}} \underbrace{\begin{bmatrix} \epsilon_1 \\ \epsilon_2 \\ \epsilon_3 \\ \epsilon_4 \\ \epsilon_5 \\ \epsilon_6 \end{bmatrix}(t)}_{\boldsymbol{\epsilon}(t)}$$

System noise distribution per time step: $\Delta\left[W_{j\in[1,6]}\right](t-u) \sim \mathrm{N}(0, t-u)$ Observation noise distribution: $\left[\epsilon_{j\in[1,6]}\right](t) \sim \mathrm{N}(0,1)$

Note: *UcorSDtoChol* converts lower tri matrix of standard deviations and unconstrained correlations to Cholesky factor,
*UcorSDtoCov* = transposed cross product of UcorSDtoChol, to give covariance, See Driver & Voelkle (2018) p11.
Individual specific notation (subscript i) only shown for subject parameter distribution – pop. means shown elsewhere.
Linearised approximation of subject parameter distribution shown.
